# A porcine bimicrobial abscess model for testing interventional treatments

**DOI:** 10.1101/2020.03.05.979088

**Authors:** Yak-Nam Wang, Andrew A. Brayman, Keith T. Chan, Keith Richmond, Wayne L. Monsky, Thomas J. Matula

**Affiliations:** Center for Industrial and Medical Ultrasound, Applied Physics Laboratory, University of Washington, Seattle, Washington 98105, USA; Vantage Radiology and Diagnostic Services, Federal Way, Washington, USA; BioDevelopment Associates, Mountlake Terrace, Washington, 98043, USA; Department of Radiology, University of Washington, Seattle, Washington 98195, USA

## Abstract

**Background:** Abscess formation is a host defense mechanism to contain the spread of infection. Abscesses can affect any part of the body and are common *sequelae* to complications of trauma, surgery, systemic infections and other disease states. Most models for abscesses are in small animals. Pursuant to the goal of developing more advanced treatments for abscesses, we sought to develop a large animal model which would reasonably mimic a fluid-filled human abscess.

**Methods:** Domestic swine were inoculated with a bimicrobial mixture of *Bacteroides fragilis* (*B. fragilis*) and *Escherichia coli* (*E. coli*) supplemented with an irritant (dextran). Inoculations were performed under ultrasound guidance in the muscle, subcutaneously or intradermally within the same animal. Fourteen days after inoculation, lesions were imaged with ultrasound, resected and prepared for histological evaluation.

**Results:** Injection of bimicrobial (aerobic and anaerobic) bacterial mixtures at multiple sites in a pig produced multiple lesions with histological features similar to encapsulated and multiloculated/multichamber abscesses often observed clinically in humans. Important salient features include the formation of a connective tissue capsule surrounding histologically nearly amorphous pus.

**Conclusions:** This paper provides the first description of a pig model for multiloculated abscesses. This animal model could potentially enable the evaluation of new technologies to replace or augment the current standard of care (image-guided percutaneous abscess drainage with antibiotics).

## Introduction

An infected abscess is defined as a focal, localized collection of infected purulent material which is delineated by a capsule composed of granulation and connective tissue [1, 2]. Breaches in physical barriers separating regional microbiomes from sterile tissue can trigger dysbiosis of both pathobionts and otherwise commensal microbes, resulting in an infection. If the infection is not resolved, the host’s immune system can wall off the infection leading to the formation of an abscess. Abscesses can affect any part of the body and are common *sequelae* of complications of trauma, surgery, systemic infections, other disease states, and drug use. Given the ubiquitous nature of bacteria and the serious threat of multi-drug resistant strains, abscesses have become a persistent global healthcare problem [3].

Aside from patient condition, treatment depends on abscess location, size and structural complexity (unilocular or multilocular). Superficial soft tissue abscesses are routinely treated with incision and drainage, which involves an incision, dissection of any loculations, debridement or drain placement. This is an extremely painful procedure which can require the use of sedation or anesthesia, particularly in young patients. Abdominal abscesses rank among the top 10 diseases for the highest 30-day rehospitalization rate [4]. Abdominal abscesses <3cm in diameter are often treated with antibiotic therapy alone [5] because indwelling catheters are physically too large to enable drainage. Image-guided percutaneous abscess drainage in conjunction with antibiotics is the preferred treatment for larger abscesses [5]. Following catheter placement, drains remain in place until there are clinical signs of abscess resolution. Patients who are not candidates for percutaneous drainage, or when percutaneous drainage is not successful (*e.g*., fails to control sepsis or fails to close fistulae), surgical intervention may be needed. Percutaneous drainage is preferred over surgical management as it is associated with higher patient survival and outcome [6]. Still, the success of percutaneous drainage depends on abscess size, location, and complexity. Multiple large, multiloculated abscesses and those containing non-drainable necrotic material or thrombus can be difficult to treat and often requires surgical intervention [7]. Risks with in-dwelling catheters include clogging, tube dislodgement, and secondary infection at the catheter access site, all of which require hospitalization and wound management. Further, the patient’s quality of life is reduced significantly by the duration of treatment with the presence of one or more in-dwelling catheters.

Given the morbidity and financial burden of percutaneous drainage, as well as the increasing problem of antibiotic resistance, the risk for secondary infection, wound management and reduced patient quality of life, new methods to treat abscesses are needed [8, 9]. As such, appropriate animal models are required in order to perform preclinical studies. Historically, five main approaches have been used to generate abscesses [10]; *viz*: 1) surgical trauma [10, 11], 2) intravenous infusion of live microbes [12], 3) fecal matter inoculation/implantation [13], 4) inoculation with a defined mono-microbial or polymicrobial load with or without an adjuvant (*e.g.* irritant, blood clot) [14, 15] or 5) a combination of the previous four [16, 17]. *B. fragilis* and *E. coli* are two common microbes used in inoculates as they are the predominant isolates in abscesses found at multiple body sites [18]. Most of these studies have been performed using small animals, and it has been challenging to develop a reliable and reproducible model which recapitulates the features of human abscesses [19]. Where evidence is presented in the primary literature, many of the reported abscesses appear to have been niduses with neutrophilic boundaries, phlegmons, or to have been pus collections constrained by folds or pockets in constitutive body membranes. There are very few animal models that have shown a collection of infected material surrounded by a capsule [9, 17, 20]. Of all these studies, there are only two studies [17, 20] that have been successful in developing true abscesses (walled off collection of infected purulent material) that are large enough to be useful in evaluation of burgeoning therapies compared with the current standard of care.

McDonald *et al.* (1980) [17] followed induced mechanical liver trauma in rabbits with an injection of *B. fragilis* (10^6^ cfu), *E. coli* (10^5^ cfu) and *Fusobacterium necrophorum* (10^6^ cfu) as either monocultures, or in combination. Injections of *E. coli* alone did not result in abscess formation and injections of *B. fragilis* alone generated small abscesses, albeit unreliably. *F. necrophorum* alone, or in combination with *E. coli* and *B. fragilis*, produced large (average diameter 4 cm) and often multiloculated abscesses. Evidence of early capsule formation was observed at 7 days, and by 28 days a well-demarcated cavity was formed from which pus could be aspirated. Despite the generation of a true abscess, this study had several limitations. First, 19% of the rabbits with abscesses died within 2 weeks of injection, and of the remainder, 22% died within 4 weeks. This limits the ability to use this model to evaluate the long-term effect of therapies. Second, *Fusobacterium ssp*. are more commonly found in abscesses of the mouth, head, neck and fingers [18], and their use could limit the significance of the study with respect to abdominal abscesses. Third, rabbits have significant anatomical and physiological differences from humans which would again limit the applicability of this model for preclinical work.

Although pigs are known for their remarkable similarities to humans, there has been only a handful of documented studies in which abscesses have been generated in pigs. Nielsen *et al.* (2009) [21] introduced an intravenous inoculation of *Staphylococcus aureus (S. aureus)* producing what the authors termed microabscesses (~20-50 μm in diameter); unfortunately, these are too small to evaluate interventional treatments. Zhang *et al.* (2013) [20] inoculated mechanically injured livers of pigs with *S. aureus*-infected venous blood/sheared clot with or without an anti- or pro-coagulant. The injections were performed directly at the trauma site or *via* the superior mesenteric vein. The largest abscesses (2 – 3 cm in diameter) were formed with the injection of an infected sheared clot in 4/5 animals. In these animals, abscesses with a complete capsule containing pus were observed 21 days after inoculation. There was no mention of the presence of multiloculi in this study, and there was a high frequency of metastatic lung abscesses generated (3/5 animals). Moreover, it is noteworthy that the cost of the pig as a study animal can be prohibitive when only generating one target abscess per animal.

In this paper, we describe the initial study in the development of a large animal abscess model in which multiple large multiloculated bimicrobial abscesses can be formed at distinct sites in the same animal. These abscesses satisfy the true definition of an abscess and are formed within a short period [1, 2]. This animal model is geared toward testing new technologies for abscess treatment and addresses some of the limitations found in other studies.

## Methods

### Culture and inoculum preparation

Prior to the day of inoculation, the dextran bead component was prepared (Cytodex-1 micro carrier beads, Sigma, St. Louis, MO USA) [22]. Briefly, a stock suspension of the dextran beads was prepared by suspending 1 g in 50 ml of EPA dilution water [23] and autoclaving. When cool, excess buffer was decanted and the remaining slurry was refrigerated until use. *E. coli* strain ATCC 25922 (both from the American Type and Culture Collection, MD, USA) and *B. fragilis* strain ATCC 23745 (American Type and Culture Collection, MD, USA) were used for all inoculations. Each species was grown separately; *E. coli* was cultured in 3% tryptic soy broth under aerobic conditions and *B. fragilis* was cultured in 3% brain heart infusion broth under anaerobic conditions. On the day of injection, *E. coli*, *B. fragilis* and sterile dextran micro carrier bead suspension were mixed in a ratio of 1:1:2 so that the final inoculum contained approximately 10^6^ cfu/ml *E. coli* and 10^6^ *cfu/ml B. fragilis,* as determined using spectrophotometric absorbance readings. The cfu number concentrations of each species as used were confirmed after the fact using Compact Dry™ EC100 and TC assay plates (Hardy Diagnostics, Santa Maria, CA, USA).

### Pig model

All procedures were approved by the Institutional Animal Care and Use Committee (IACUC) at R&R Rabbitry (Stanwood, WA). Three female domestic swine (Yorkshire/Hampshire cross, age 3-6 months, weight 45-55 kg) were used in this study. On the day of inoculation, animals were sedated (Telazol/Ketamine/Xylazine) and maintained under a surgical plane of anesthesia with isoflurane. All animals were instrumented to monitor heart rate, ECG and blood oxygen saturation and temperature. The haunches and abdomen of the animal were depilated, cleaned and sterilized with serial rounds of isopropyl alcohol. Injections were performed under ultrasound guidance into 6 separate sites in a single animal: 1.5 cm deep into the left and right biceps femoris (6 injections total for all animals), and either subcutaneously (7 injections total) or intradermally (5 injections total) on either side of the midline. For each animal, 5 or 10 ml of the inoculate were injected at each site (a total of 3 sites with 5 ml, 15 sites with 10 ml) and the location was marked with dye. The injection volumes were chosen as our preliminary studies (data not presented) have shown that injections of 2 ml were unreliable at generating lesions, and when formed, these were small (~1cm in diameter). After the inoculations, the animals were recovered. Upon recovery post injection, the animals were observed for signs of distress. The animal and injection sites were monitored every day for signs of abscess growth and clinical signs. Animals were observed for signs of pain and distress, reduction in appetite, loss of weight and mobility. The observational outcomes are described in Results.

Two weeks after the inoculation, the animals were sedated and anesthetized. The injections sites were palpated and imaged with ultrasound. An 18G needle was used to try to determine if some of the abscess contents were liquid enough to aspirate. When a small sample of the contents could be aspirated, this was placed into culture and plated on the Compact Dry™ EC100 and TC plates. The animal was euthanized and all the lesions were dissected out *en bloc* as much as possible. If lesions were bisected, the lesions were evaluated grossly. Lesions were fixed in 10% neutral buffered formalin to be processed for histological evaluation.

### Histopathology

Fixed lesions were grossed and processed to maintain the lesion as whole crosssections of the lesions as much as possible. Samples were sectioned and stained with hematoxylin and eosin (H&E) for general tissue morphology, Masson’s trichrome stain to visualize the presence of a fibrous capsule, or Gram stain to evaluate the presence of bacteria. Lesions were evaluated for the presence of a connective tissue capsule; the extent of the capsule, if present; the presence of infected material, cell debris and inflammatory cells; and the presence of loculations.

## Results

All animals tolerated having 5 or 10 ml of a bimicrobial mixture injected into multiple sites. During the injection, the generation of a volume within the tissue could be observed by ultrasound guidance. This cavity remained for many of the injections during the course of the anesthetic period. Immediately after injection, some sites did not exhibit any change in gross appearance, but some sites exhibited a large bulla, mainly with the intradermal injections. On recovery, and during the two weeks after recovery, none of the animals were febrile or displayed signs of pain. Lesion generation could not be grossly detected at all the sites over the two-week period during the daily monitoring and none of the lesions broke through the surface of the skin.

### Intramuscular Injections

As mentioned earlier, there were one 5 ml injection and five 10 ml intramuscular injection. At necropsy (two weeks after inoculation), none of the intramuscular lesions were palpable. There was no skin discoloration above the injection site (Figure 1). However, lesions were clearly seen with ultrasound imaging (Figure1). There were no differences between the 5 ml and 10 ml injections detectable using ultrasound imaging. There was some variation in the hyperechoity of the lesions, but no pattern could be discerned. In the site injected with 5 ml, the lesion was accidentally cut through during tissue harvest revealing liquid pus which readily flowed out of the cavity (Figure1). However, not all lesion contents could be aspirated using an 18G needle.

**Figure 1.**
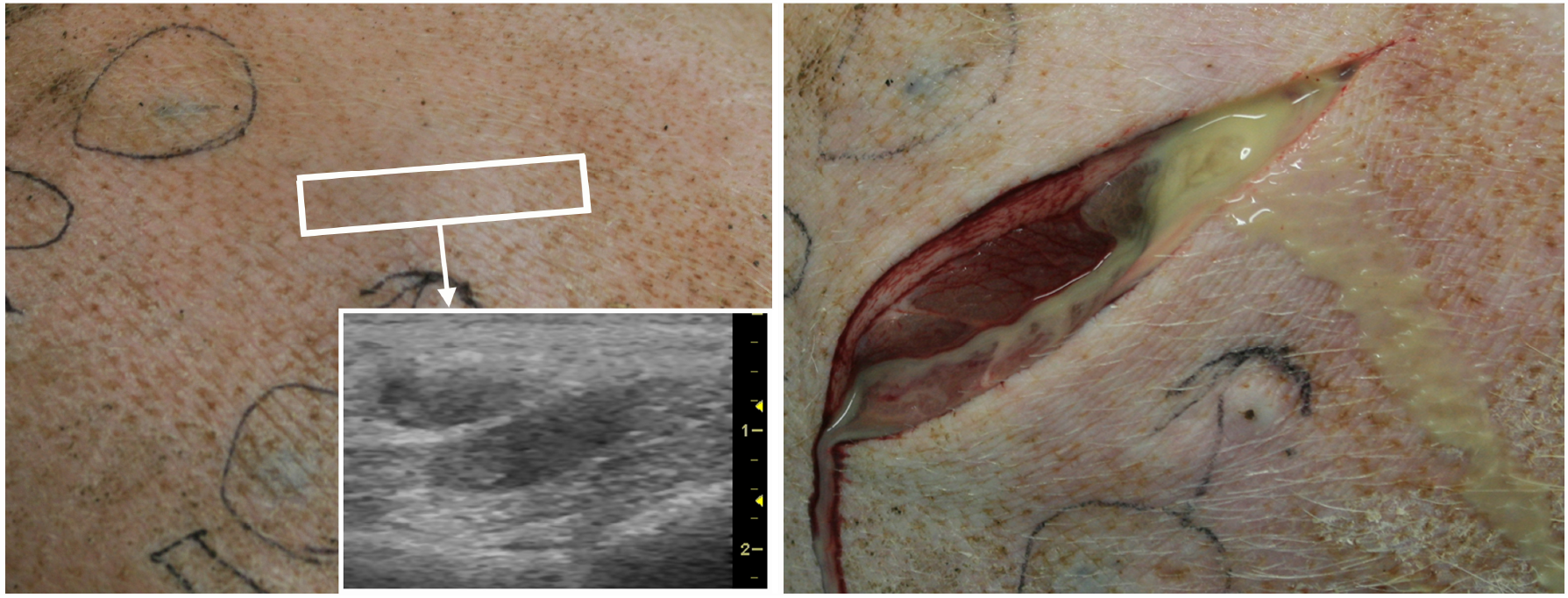
Photo of skin surface (left) two weeks after inoculation. Ultrasound image (insert) taken from a region over the abscess (white rectangle) shows two distinct hypoechoic cavities within the muscle. Scalar values are in cm. On necropsy, the collection was breached and a purulent yellow liquid flowed out of the cavity (right).

Histological evaluation revealed that all injections (1/1 of the 5 ml injection and 5/5 of the 10 ml injections) generated lesions with well-defined connective tissue capsules with granulation tissue and multiple loculations within many of the chambers (Figure 2). The loculations were separated by dense connective tissue. The cavity contained aggregations of neutrophils within a field of proteinaceous and cellular debris. The Gram-stain sections revealed Gram negative-bacteria within the capsule (data not shown). In some of the lesions, fragments of muscle tissue could also be seen. Dextran was present in all the lesions.

**Figure 2.**
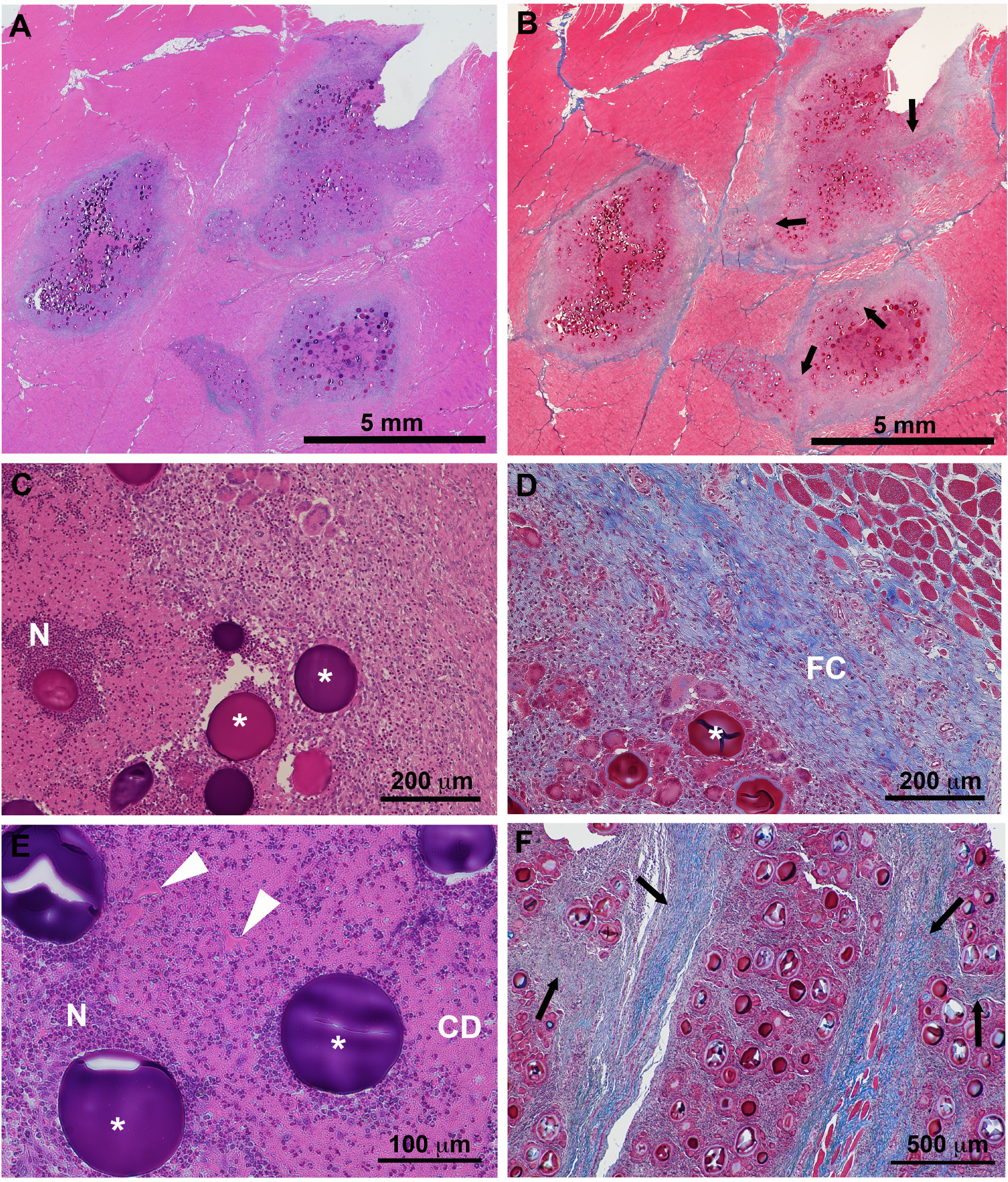
Hematoxyin and eosin (left column) and Masson’s trichrome (right column) stained sections of intramuscular lesions at various magnifications. Multiple chambers (panels A and B) of infected material surrounded by a fibrous connective tissue capsule (FC in panel D) were observed at all sites. Loculations separated by connective tissue septae (black arrows in panels B and F) were observed within the chambers. Magnification of H&E sections (panels C and E) revealed cavities containing aggregations of neutrophils (N) and inflammatory cells within a field of cellular debris (CD). Some amorphous proteinaceous material (white arrow head) was also observed in many of the cavities. Dextran (asterisks) was present in all lesions.

### Subcutaneous Injections

At necropsy, the majority of the subcutaneous lesions could be seen grossly and were palpable. There was no skin discoloration above the injection site and no perceptible increase in skin temperature. The majority of lesions could be clearly observed with ultrasound imaging. Almost all of the injections resulted in the generation of a lesion (2/2 of the 5ml injections and 4/5 of the 10 ml injections). All but one of these lesions had a well-defined collagenous capsule. In the one lesion in which no defined capsule was present (Figure 3), there was a large region of structural distortion and inflammation. In the center, there appeared to be a dense inflammatory response with a wide rim of distorted tissue caused by the apparent thickening of the elastic septa and the presence of large vacuoles (Figure 3).

**Figure 3.**
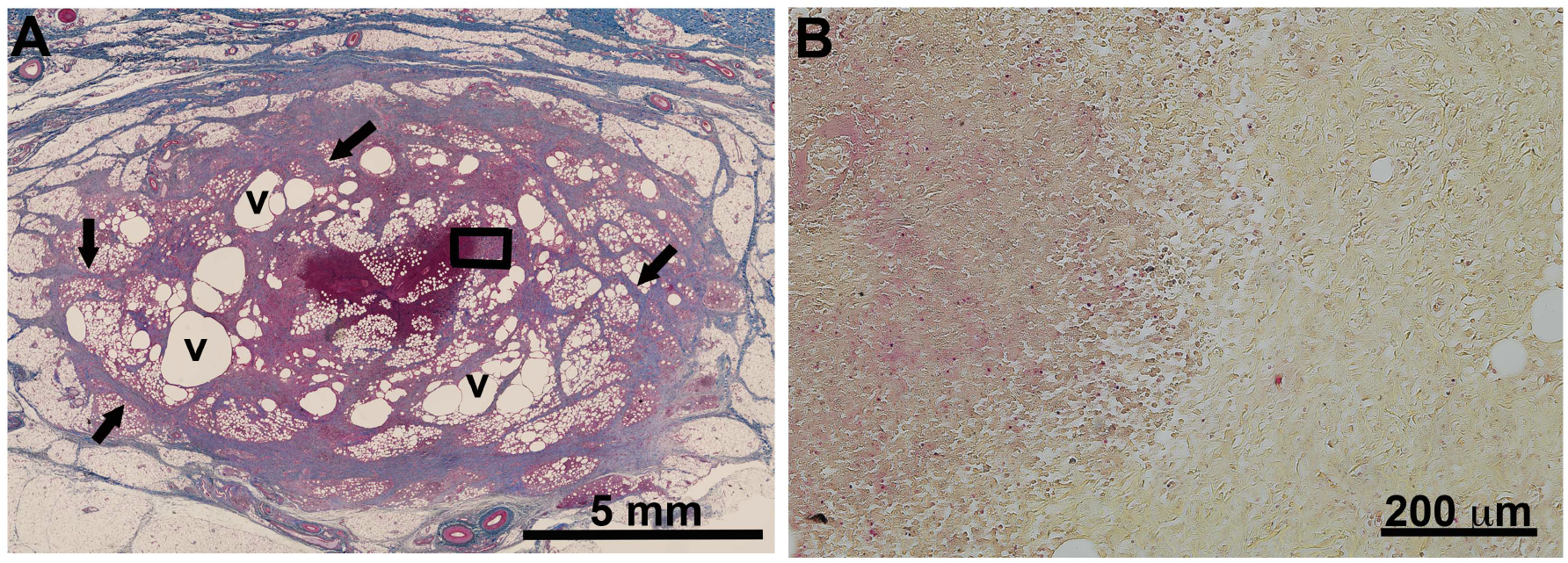
Masson’s trichrome stained lesion (panel A) without a defined capsule. The lesion contained a focal center of inflammation surrounded by a ring of distorted tissue with thickened septae (black arrows) and vacuoles (v). A magnified image (panel B) of the focal inflammatory center (box in A) in a Gram stained section showed Gram-negative bacteria (pink staining) concentrated within the inflammatory center and not in the surrounding tissue.

### Intradermal Injections

Recall that there were two 5 ml and five 10 ml SQ injections. Immediately after injection, a distinct swelling was apparent at all intradermal injection sites, and at many of the larger injection sites, the skin was reddened. In some instances the inoculum may have extended into the subcutis. Despite the extent of the swelling at the site of some of the lesions, none of the lesions perforated the skin during the two-week follow-up period. At necropsy, 4/5 of the 10 ml injection sites could still be seen as distinct nodular foci, broadly similar in appearance to that seen immediately post injection. Only half of the lesions contained material which could be aspirated *via* an 18G needle. However, upon ultrasound imaging, more than one distinct cavity could be seen in all of the lesions (Figure 4). These cavities were similar in appearance, and were characterized by a hyperechoic nucleus within a slightly less hyperechoic cavity (Figure 4). A well-defined capsule was present around all the lesions (Figure 5). Each cavity contained aggregations of the neutrophils admixed with proteinaceous and cellular debris. The Gram stain revealed the presence of Gram-negative bacteria within all cavities; in some lesions, these were observed to be lining the inner wall of the cavity. As with the intramuscular lesions, inflammation was also observed as part of the capsule. Dextran was present in all lesions. The one lesion that could not be seen as a distinct nodular focus-appeared on ultrasound imaging as a region where the tissue seemed to be disrupted (Figure 5). Histologically this lesion appeared very similar to the diffuse lesion that was generated by of the subcutaneous injections (Figure 3) and as such represents more a condition of cellulitis.

**Figure 4.**
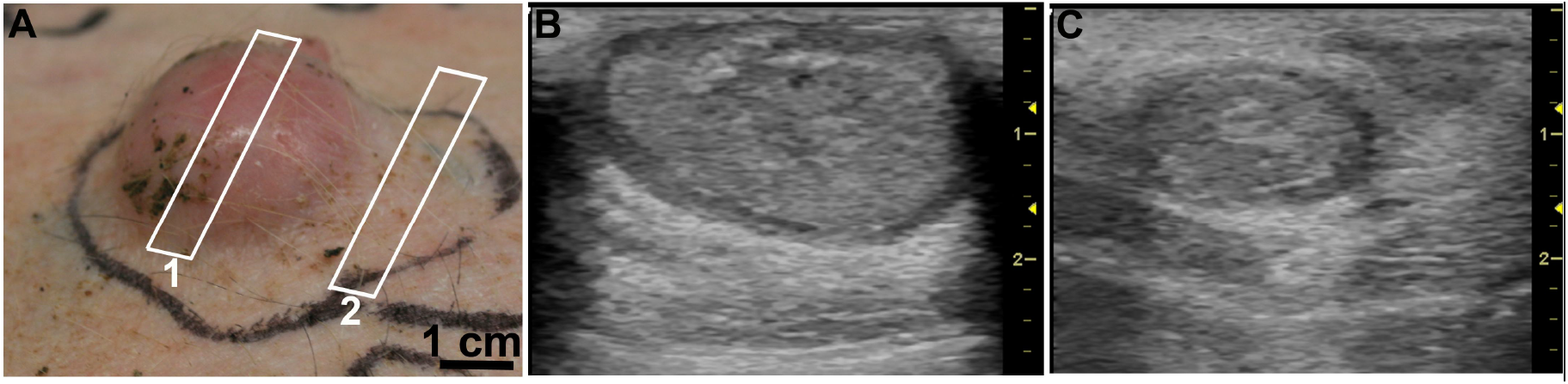
Photo (panel A) of an intradermal lesion 14 days post inoculation. Ultrasound imaging revealed multiple chambers when scanning over the lesion. Ultrasound images B and C were taken with the ultrasound probe placed at site 1 or 2 respectively. Scale on right of ultrasound images are in cm.

**Figure 5.**
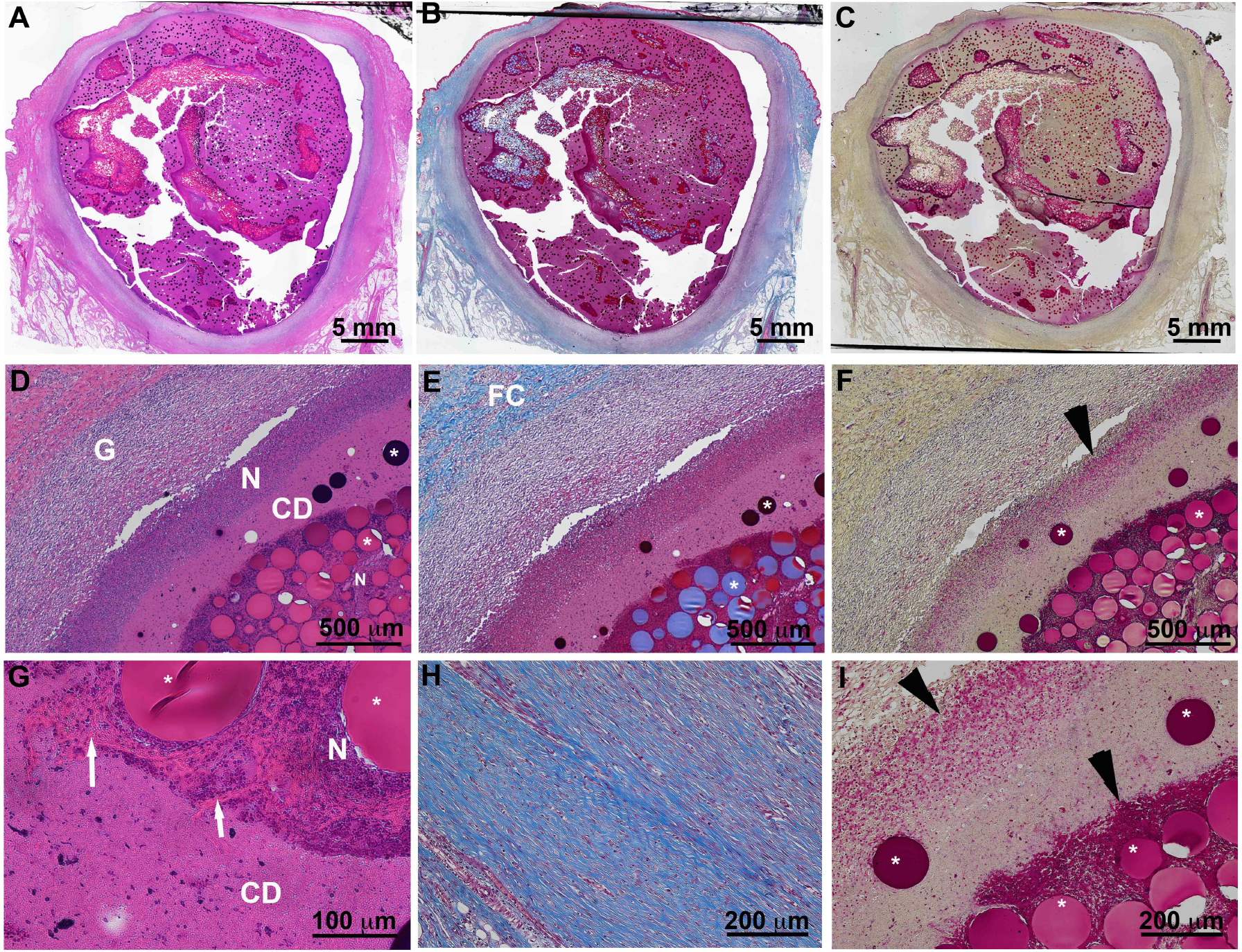
Hematoxylin and eosin (left column), Masson’s trichrome (middle column) and Gram (right column) stained intradermal lesions at various magnifications. Panels A, B and C show a cross-section through a large lesion. Panels D-F and G-I present views at progressively greater magnifications in the same sctinos and staining regimens. All lesions had a thick and robust fibrous connective tissue capsule (FC) which contained regions with neutrophil aggregates (N) containing phagocytized bacteria (black arrowhead) and inflammatory cells in a sea of cellular debris (CD). Within the neutrophil aggregates there were amorphous, eosinophilic matter (white arrow). Around the rim of the lesion there was (black arrowhead) neutrophils (N) and beyond that, a ring of granulation tissue (G).

## Discussion

Antibiotic use is ubiquitous for the treatment of all kinds of infections, irrespective of the location, size, or type. Aside from the unpleasant side effect of changing the normal microbial flora, antibiotic resistance has been called the most significant threat to modern medicine by the World Health Organization (WHO) [3]. New technologies and drugs are therefore desperately needed to treat infections such as abscesses. An appropriate animal model which significantly recapitulates the human condition is needed to test such technologies and drugs. Although small animals enable large cohort studies, they have significant anatomical and physiological differences from humans which would limit the significance of such studies. Moreover, the abscesses generated in these models are often unreliable and not always reproducible, and not similar in structure to human abscesses.

We here present an animal model in which bimicrobial multiloculated abscesses which capture salient features of human abscesses can be reproducibly formed in a pig. Pigs are known for their remarkable similarities (anatomical, physiological and immunological) to humans [24, 25] and have become one of the dominant animal species used in translational research. Out of respect of the 3R principles (replacement, reduction and refinement) for animal use, we looked to the porcine wound healing model in which multiple wounds are generated in the same animal.

The abscesses in this model were located in the muscle, skin or subcutaneous regions. Of the 18 injections made, we successfully generated 16 abscesses which satisfied the histopathological description of an abscess: a focal, localized collection of purulent material delineated by a wall of connective and granulation tissues [1, 2]. There are currently two papers published describing the generation of abscesses in a large animal model; one using rabbits [17] and the other minipigs [20]. Both models successfully generated a relatively large abscess. However, there were significant limitations of each model including the high mortality rate in rabbits, lack of multiloculation, the generation of only one target abscess per animal and the type of bacteria used. The current model overcomes the limitations of these studies.

When abscesses were formed in this model, they were all multiloculated, with loculi separated by collagenous septae. Multiple loculations are a characteristic of complex human abscesses, complicating the ability to control infection and adequately drain the collection. They often require multiple drain placements and, in some cases, surgery is needed. This salient feature is essential to recapitulate in an animal model to better represent the more challenging cases. In this work, the larger chambers were easily visualized using ultrasound imaging, a modality commonly used to guide percutaneous drainage.

Abscesses described here were generated using a bimicrobial mixture of *E. coli* and *B. fragilis*; the predominant isolates in abscesses found at multiple body sites [18]. *E. coli* is the most prevalent Gram-negative pathogen in humans so the consequence of antibiotic resistance would be particularly concerning. The rates of antibiotic resistance are rapidly rising. Multi-drug resistance in *E.coli* increased from 7.2% during the 1950s to 63.3% during the 2000s [26]. Not only is *B. fragilis* the most virulent *Bacteroides* species, but it has been reported to have the highest resistance rates of all anaerobic pathogens [27]. The treatment of abscesses generated with these pathogens could, therefore, hold great clinical significance.

One main limitation of this study is the inclusion of dextran in the inoculant. Dextran was used as an irritant to elicit a persistent inflammatory response. It is not known how or if the presence of the dextran will alter the treatment or drug being tested on the abscesses generated in this model. It is possible that the injection of a large volume of the bimicrobial mixture would be enough trauma to elicit a robust and sustained inflammatory response and the subsequent generation of an abscess. However, this was not evaluated since many studies have reported the need for an irritant [14, 15, 28, 29]. It is also possible that pre-injuring the tissue before inoculation might also eliminate the need for the dextran; this will be evaluated in future studies.

These lesions were only followed out to two weeks post-injection. During the two weeks, none of the animals showed any signs of distress from the injections nor the presence of the lesions as they developed. Animals were not febrile, and none of the lesions erupted through the skin surface. Future studies will expand the time after inoculation to determine how the lesions evolve over longer periods so that the utility of this model for acute as well as chronic studies can be assessed.

## Conclusion

This is the first description of a pig model for multiloculated abscesses, a difficult and costly to treat complication from infection. This animal model could potentially enable the evaluation of new technologies to replace or augment the current standard-of-care (image-guided percutaneous abscess drainage with antibiotics).

## Acknowledgements

The authors acknowledge and appreciate, with sincere thanks, Denny Liggitt, Department of Comparative Medicine, University of Washington, for the many discussions and careful reading of this manuscript.

